# Rare RNA Polymerase II failure modes mark the cancer-driving genes most affected by epigenetic perturbation

**DOI:** 10.64898/2026.05.28.728581

**Authors:** Yaw Asante, Berkley Gryder

**Affiliations:** Department of Nutrition, Case Western Reserve University, Cleveland, OH, USA; Department of Genetics and Genome Sciences, Case Western Reserve University, Cleveland, OH, USA

**Author notes:** Corresponding authors: Yaw Asante, Berkley Gryder.

## Abstract

RNA Polymerase II (Pol2) transcribes genes through a complex life cycle (initiation, pausing, elongation, co-transcriptional splicing, termination, and recycling). Chromatin immunoprecipitation of Pol2 before and after chemical perturbation has identified promoter-proximal accumulation (pausing) as a critical step in the transcription genome-wide. However, the full landscape of Pol2 responses has not been well characterized. Here, we introduce a tool for comparing Pol2 Activity State Shifts (compPASS), a computational pipeline which uses data from paired ChIP-based approaches to assign genes to one of eight distinct modes by Pol2 response under different forms of perturbation. In multiple cancer types and drug contexts, we show that compPASS identifies previously undescribed Pol2 failure modes with important implications for gene regulation. By looking past pausing, compPASS exposes Pol2 failure modes (clogging, entry, gain, loss) that are rare but pinpoint the genes most relevant to cancer cell state changes in response to therapy, turning a single paired Pol2 ChIP-seq into a mechanistic map of shifting transcriptional states.

## INTRODUCTION

RNA Polymerase II (Pol2) transcription in eukaryotes is regulated at distinct stages by various chromatin-associated proteins. General transcription factors and the pre-initiation complex are essential for the recruitment of Pol2 to gene promoter sites.Canonical interactions between the complexes DSIF and NELF, followed subsequently by p-TEFb, mediate the arrest and progression of Pol2 respectively^1^. However, other genes like immediate early genes, growth factors and even lineage-specific transcription factors can play a downstream role in mediating Pol2 accumulation upstream of the gene (pausing) or, conversely, enabling Pol2 movement into the gene body (releasing) under specific conditions^2^. The orchestration of recruitment, initiation, pausing, release, elongation, termination and recycling of Pol2 have been identified and characterized through both fluorescent microscopy and functional genomic perturbation of putative epigenetic regulators^2,3^. Each of these steps is driven by proteins which can be targeted to disrupt the efficacy of transcription.

A common way of measuring changes in Pol2 activity at gene loci is through chromatin immunoprecipitation and sequencing (ChIP-seq). Aligning the chromatin reads associated with Pol2 provides us with a continuous ledger of the amount of Pol2 along the entire human genome, with gene proximal sites being the most enriched for total signal. The profile of this data at gene loci is typically measured at distinct regions of interest relative to the transcription start site (TSS) and the transcription end site (TES) which, compared to all other genes, inform us of how well Pol2 is processing transcripts at that site. While the presence of Pol2 along the gene has been shown to correlate with active expression in some cell types^4^, Pol2 at the TSS site is often a key focus.

Typically, the density of reads near the TSS (sometimes called the promoter-proximal region) is compared to the density of reads along the gene body. This has been called the pausing index or pausing ratio and the cumulative distribution of these values between untreated and perturbed cases has been widely used to show the impact of a drug or gene knockdown on transcription^2^. Induction or ablation of pausing has been shown to be transcriptionally relevant in many cell types^5,6^. Further still, pausing reflects distinct epigenetic contexts such that the same perturbation may impact similar length genes in very dissimilar ways, based on how many enhancer regions support their expression^7^.

Measuring Pol2 pausing is not the only way that we can use Pol2 activity shifts to identify biologically relevant gene perturbation. A previous study^8^ quantified the ratio of reads between a region upstream of the TSS (here, called the promoter region) and the TSS region following small molecule treatment to measure Pol2 Stalling Index. In that case, MEK signaling prevented Pol2 entry into the pause site at *MYOG*, which was shown to be crucial in preventing differentiation and maintaining cancer state in fusion-negative rhabdomyosarcoma. Additionally, our group demonstrated in prior work^9,10^ a novel concurrence between reduced gene expression and Pol2 accumulation in a small but crucial set of genes in fusion-positive rhabdomyosarcoma. Previously identified master and core-regulatory transcription factors were abundant among the genes which experienced the greatest change in the ratio of number of reads upstream of the TES relative to the number of reads several thousand base-pairs downstream of the TES, a phenomenon which we then measured as the unloading ratio. Each of these metrics captures in a snapshot the functionality of Pol2 at stages of transcription, but these modes have not previously been examined altogether.

Given that Pol2 profiles, as measured through ChIP-seq and HiChIP, change in ways that are reflective of important biological events^5^, we developed the comparing Pol2 Activity State Shifts (compPASS) pipeline to measure these ratios and assign response classes to each gene genome-wide. Additionally, given the lack of a formalized approach to the study of Pol2 activity shifts, we outline a set of definitions for regions of interest and ratios for future study. This framework classifies every Pol 2-perturbed gene into one of eight response modes from a single paired ChIP-seq experiment. Across CDK9, MEK, and p300/CBP perturbations in three cancer contexts, compPASS reveals that the rarest modes, rather than the most abundant modes, are often the ones most relevant to disease-driving or phenotype-relevant genes.

## RESULTS

### Analytical framework, use-cases and workflow of compPASS pipeline

RNA Pol2 drives gene transcription in discrete stages which correspond to regions of the gene (Fig. 1A). However, it does not occupy these stages in equal measure. Pol2 accumulates at the very start of genes (the transcription start site or TSS) and its movement through discrete regions of a gene is coordinated by cofactors that, if perturbed, can modulate Pol2’s gene body distribution. ChIP-seq on Pol2 provides a genome-wide snapshot of activity at the moment of crosslinking and captures where along the body of each gene Pol2 is binding. The more Pol2 at a given site, the more reads captured in ChIP-seq and alignment can reveal Pol2 positioning across tens of thousands of genes down to the specificity of a few base pairs. Neither this level of precision nor this scale is achievable by other methods of studying transcription such as microscopy.

**Figure 1:**
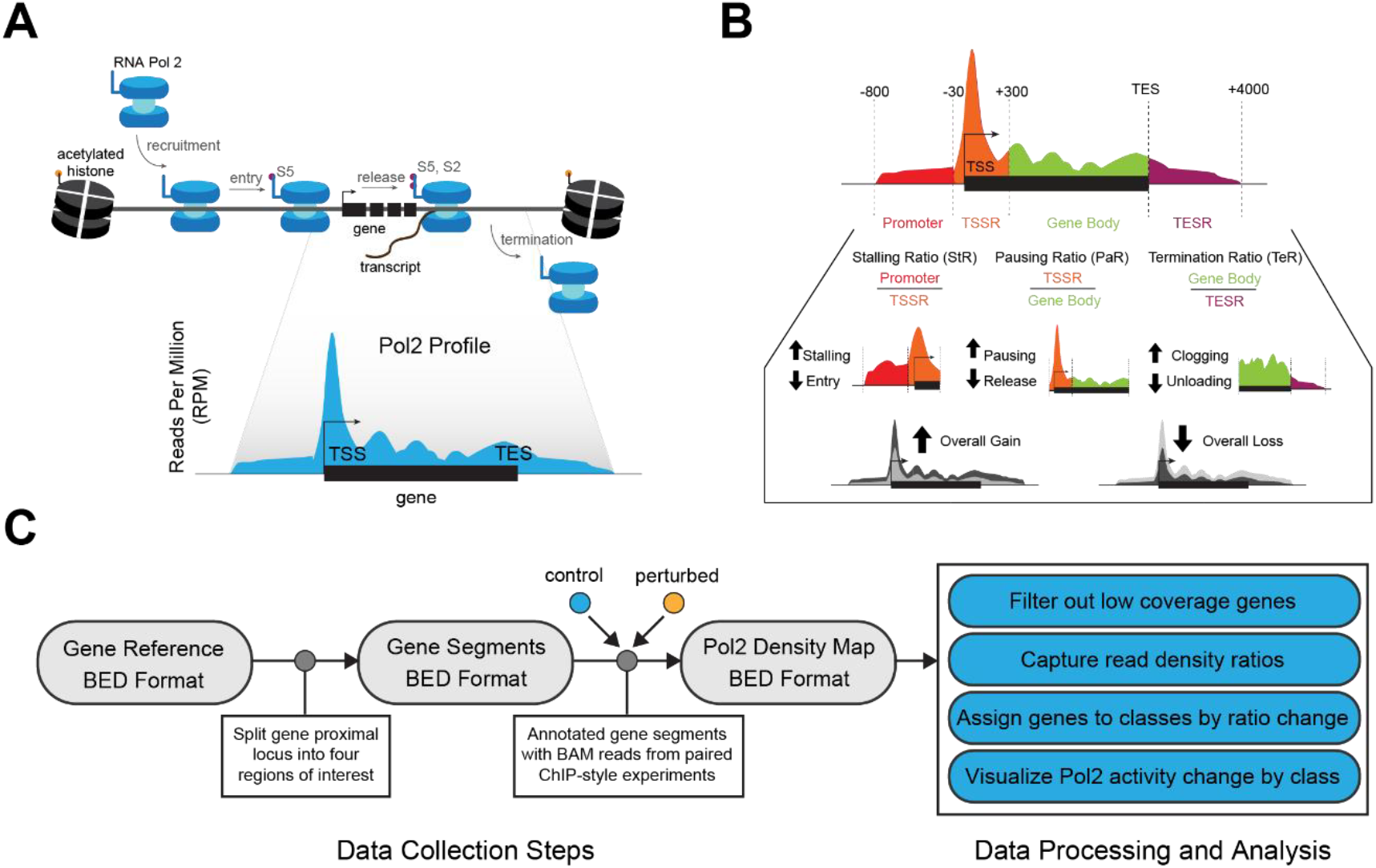
The compPASS pipeline uses ChIP-type data to capture modes of Pol2 response to perturbation. **A**. Diagram of RNA Polymerase II stages (top) and how Pol2 positioning appears in ChIP-style data in genome-browser view (bottom). **B**. Diagram of the four gene-proximal regions used to capture Pol2 read coverage, calculate the three ratios and assign the eight final states **C**. Workflow of compPASS program with two paired ChIP-style inputs.

The sequential steps of Pol2 movement along the gene proximal site are aggregated into a Pol2 profile that reflects the relative abundance of Pol2 along the gene (Fig. 1A). Several ChIP-style approaches capture Pol2 positioning genome-wide (Supplementary Fig. 1A). Although more RNA Pol2 often means more transcription of that gene, there are genes with abundant Pol2 that fail to produce complete transcripts, as Pol2 has several sequential biochemical steps that may bottleneck a gene’s expression (Supplementary Fig. 1B).

After initiation, Pol2 typically transcribes 30-100 base pairs past the TSS and halts at the “pause site” until additional co-factors attach the required marks to push it into elongation. It is standard to capture the extent of this promoter-proximal pause at a given gene by calculating its pause ratio. The pause ratio is the ratio of read density at the TSS proximal region over the read density in the gene body. A high pause ratio reflects that much more Pol2 is seated at the starting line rather than advancing along the gene and productively producing transcripts. An increased pause ratio following perturbation shows that the gene is Pausing, meaning that Pol2 is failing to progress into the gene body. The inverse, where more Pol2 is going into the gene, is called Releasing, when more Pol2 is released to actively elongate. While the pause ratio gives us a measure of how well Pol2 is performing at two sites important for orchestrated transcriptional control, it does not reflect the full breadth of the Pol2 lifecycle. We extend the logic and mathematics of Pausing further and here consider the region upstream of the TSS and the region downstream of the TES to measure changes in additional biochemical steps of transcription.

The Stalling Ratio (equal to the Stalling Index^8^) compares the read density at the promoter (-800 bp to -30 bp from the TSS) to the read density at the TSS/pause site (-30 bp to +300 bp). This captures the amount of Pol2 successfully arriving at the TSS region, with increases in the Stalling Ratio reflecting a failure to enter the pause site. Genes with a substantive increase in Stalling Ratio are classified as *Stalling*, while genes with the inverse are classified as *Entering*.

We define the Termination Ratio as the ratio of read density at the gene body (from TSS +300 bp to the TES) over the read density of Pol2 binding in a 4kb window downstream of the TES. These values reflect whether Pol2 is progressing through the Poly-A/cleavage process properly or experiencing termination failure. Major increases in Termination Ratio after perturbation marks a gene classified as *Clogging* (failing to move past the TES), whereas major decreases mark a gene classified as *Unloading* (Fig. 1B).

Two additional states, for genes lacking any ratio changes, are identified when Pol2 undergoes a balanced overall gain (*Pol2 Gain*) or loss (*Pol2 Loss*) across all regions. Many genes usually experience no significant mode change.

The compPASS tool applies this framework across a provided subset of genes. It annotates segmented gene regions with Pol2 ChIP-seq/HiChIP read counts in two conditions, assigns each one of eight Pol2 activity state changes defined above, and tests predefined gene groups for enrichment of the major failure modes (Fig. 1C).Genes very rarely exhibit Pol2 shifts in more than one of these modes. compPASS provides results such that users can interrogate commonalities underlying types of state changes with biologically relevant implications.

### compPASS captures a diversity of Pol2 changes across treatments

Previous studies which employed Pol2 pausing analysis did so generally to highlight the cumulative change in pausing ratio between known, biologically relevant sets of genes. Rather than observing changes among known genes, we seek to first classify sets of genes de novo from their ratio change. This requires that we clarify the difference between genes which see changes in their Stalling, Pausing and Termination ratios and genes which are truly *Stalling, Pausing* and *Clogging* based on non-trivial changes in both the ratio of interest and the fold-change of read density at regions of interest. We require four conditions for classification into each of the six ratio-shift modes. For example, for a gene to be labeled as *Pausing*, it must show a substantial increase in TSS region Pol2 reads, a loss in gene body region Pol2 reads, a substantial increase in Pausing Ratio, and a Pause Ratio >= 2 (Supplementary Fig. 2A).

**Figure 2:**
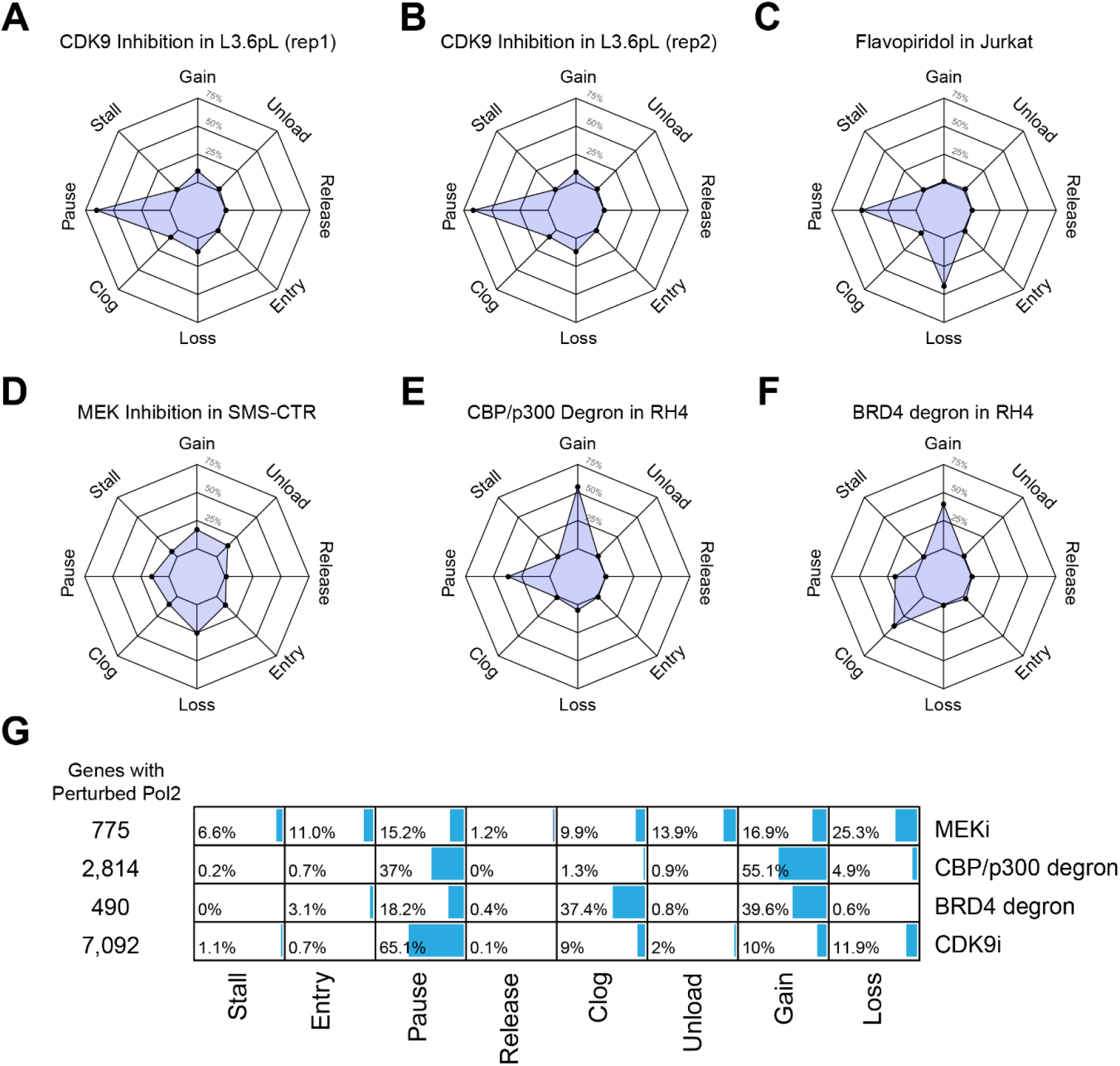
Chemical perturbation causes multiple infrequent, but substantial shifts in Pol2 activity across cancers. **A**. – **F**. Radar plots showing the percentage of perturbed genes in each of the eight final states across disease contexts and chemical perturbations. **G**. Bar plots showing the percentage of perturbed genes in each final state and the total count of perturbed genes.

We used publicly available biological triplicate Pol2 ChIP-seq in pancreatic adenocarcinoma cells (L3.6pl) to evaluate the expected variation in the amount of Pol2 identified across promoter, TSS, gene body and TES regions of genes. For a set of over 14,000 genes which have non-trivial amounts of Pol2 across the three replicates of the basal state (DMSO-treated), we calculated the per-gene variability in Pol2 abundance. We calculate variability as the log2 of the max read count over the minimum read count and plotted the distribution. We opted to use the approximate 90^th^ percentile value as the cutoff for change at a given region to filter out cases of Pol2 read change that could be typical expected deviation between Pol2 ChIP experiments (Supplementary Fig. 2B). The same concept was used to identify the change of ratios themselves, with all three ratios having a 90^th^ percentile value around 1, here reflecting the expected log2 fold change of ratios between basal and perturbed state (Supplementary Fig. 2C).

A key output of the compPASS pipeline is a radar plot showing the percentage abundance of perturbed genes in each category. CDK9 inhibition interferes with the function of p-TEFb and promotes release failure. Following the specific inhibition of CDK9, we see that most perturbed genes are *Pausing* in L3.6pL cells across two replicates (Fig. 2A, Fig. 2B). In the more complex case of flavopiridol treatment, which causes pan-CDK inhibition, we see an abundance of *Pausing* but also of overall Pol2 loss (Fig. 2C). Small molecules which induce more severe changes in cell state, like MEK inhibition (Fig. 2D), degradation of the co-activators EP300 and CREBBP (Fig. 2E) and degradation of p-TEFb co-factor and epigenetic reader BRD4 (Fig. 2F), show acutely different radar plot profiles. These variations in perturbed gene final states, as well as the number of substantially changed Pol2 profiles, demonstrate the value of looking beyond pause ratio. Most perturbed genes, in all but the CDK9i case, experience something other than pausing (Fig. 2G).

### compPASS highlights Pol2 failure modes beyond pausing after CDK9i

In pancreatic adenocarcinoma (PDAC) cells, enitociclib causes cell death through the inhibition of the CDK9. CDK9 is a core component of the p-TEFb complex and, without the ability to phosphorylate Pol2 at its C-terminal domain, transcription is unable to progress past the pause site. Broadly, we can term the expected mechanism of action for the drug as release failure (Fig. 3A).

**Figure 3:**
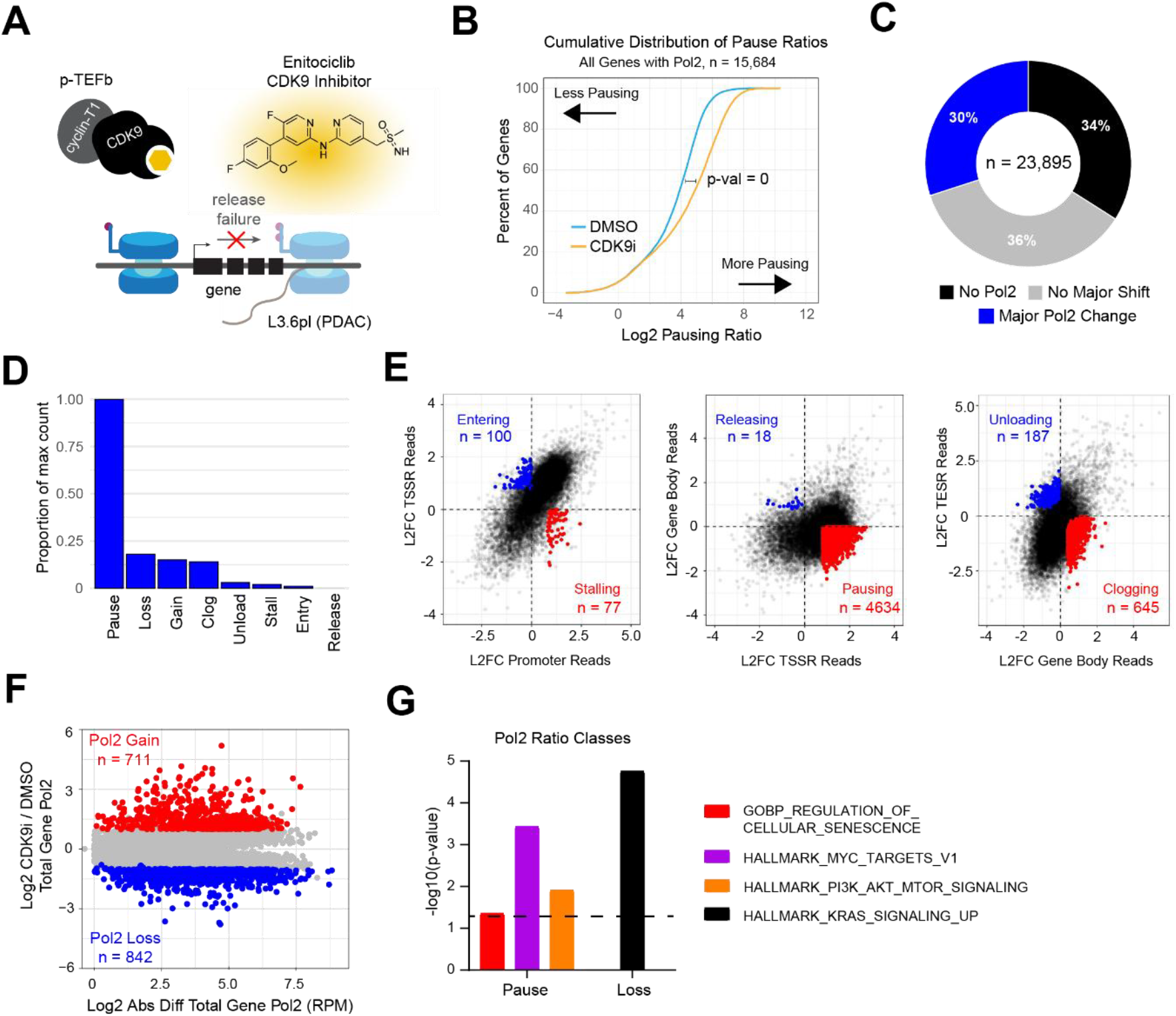
CDK9 inhibition causes both canonical pausing and distinct Pol2 loss events at cancer critical genes. **A**. Model of enitociclib action at genes in L3.6pL pancreatic adenocarcinoma cells. **B**. Cumulative distribution plot of filtered genes between DMSO and CDK9i treated states. **C**. Donut plot showing percentage of genes showing major Pol2 perturbation, no major change and not considered due to low Pol2. **D**. Bar plots showing the proportion of perturbed genes in each Pol2 state relative to the most frequent state. **E**. Scatterplots showing genes in each state based on the distribution of fold change values from their respective gene-proximal regions. Pol2 states reflecting a gain in the preceding region (Stalling, Pausing, Clogging) in red, Pol2 states reflecting a loss in the preceding region in blue. **F**. Scatterplot showing the log2 absolute difference of total Pol2 against the fold change of total Pol2. Total Pol2 is the sum of Pol2 at all gene-proximal regions. Pol2 gain genes in red, Pol2 loss genes in blue. **G**. Bar plot showing the enrichment of genes in Pol2 final states for known functional sets of genes. Dashed line shows the corresponding cutoff for p values < 0.05.

To capture substantive changes, we first filtered out 8,211 genes due to low Pol2 and reviewed the Pol2 landscape changes of 15,684 genes between control and drug treated cases (Supplementary Fig. 3A). A conventional cumulative distribution analysis of pause ratios across genes shows a highly significant shift of genes towards pausing (Fig. 3B). Applying compPASS methodology, we see that only 30% of the original gene set, representing almost 45% of currently considered genes, show a major Pol2 shift (Fig. 3C). Of the majorly perturbed set, most show *Pausing*, consistent with the release failure expectation (Fig. 3D) and the *Pausing* profile is strong across over 4,600 genes (Supplementary Fig. 3B). However, there are substantial subgroups of genes experiencing general *Pol2 Gain, Clogging* and, as the second-most abundant mode, *Pol2 Loss* (Fig. 3E, Fig. 3F). These Pol2 states reflect distinct profiles relative to standard *Pausing*. A hypergeometric test for enrichment across all Pol2 states shows most lack strong enrichments (Supplementary Fig. 3C). *Pol2 Loss* genes are enriched to a highly significant degree for KRAS signaling up genes. *Pausing* genes are also enriched for MYC targets, MTOR signaling genes and cellular senescence genes (Fig. 3G). Some negative regulators of cell senescence like *TERF2* show strong *Pausing*, suggesting inactivation upon enitociclib treatment and consistent with known anti-cancer impacts of this treatment in PDAC cells (Supplementary Fig. 3D).

### compPASS clarifies the impact of Pol2 entry on differentiation in FN-RMS

*MYOG* is a key regulator of myogenesis, the process of epigenetic reprogramming to fully commit progenitor cells to form muscle fibers. During myogenesis in healthy/normal cells and tissues, *MYOG* transcriptional activation is achieved by control elements around the gene (enhancers) which gain large amounts of histone acetylation at H3K27ac, a mark of active enhancers. In Yohe et al^8^, they find that mutant RAS signal causes an epigenetic block of differentiation in RAS-driven rhabdomyosarcomas. Upon inhibition of the RAS signal (MEK or ERK inhibition), *MYOG* is strongly activated at the transcriptional level. While *MYOG* goes on to completely remodel the epigenome and differentiates the tumors into muscle-like cells, the epigenetics (histone modification patterns) at MYOG itself remain unchanged. This mystery prompted a closer mechanistic evaluation of Pol2 at *MYOG*. Looking at RNA Pol2, the authors proposed a mechanism of action by which trametinib, an MEK inhibitor, caused *Entry* at *MYOG*, causing fusion-negative rhabdomyosarcoma cells to exit their undifferentiated state (Fig. 4A, Fig. 4B).

**Figure 4:**
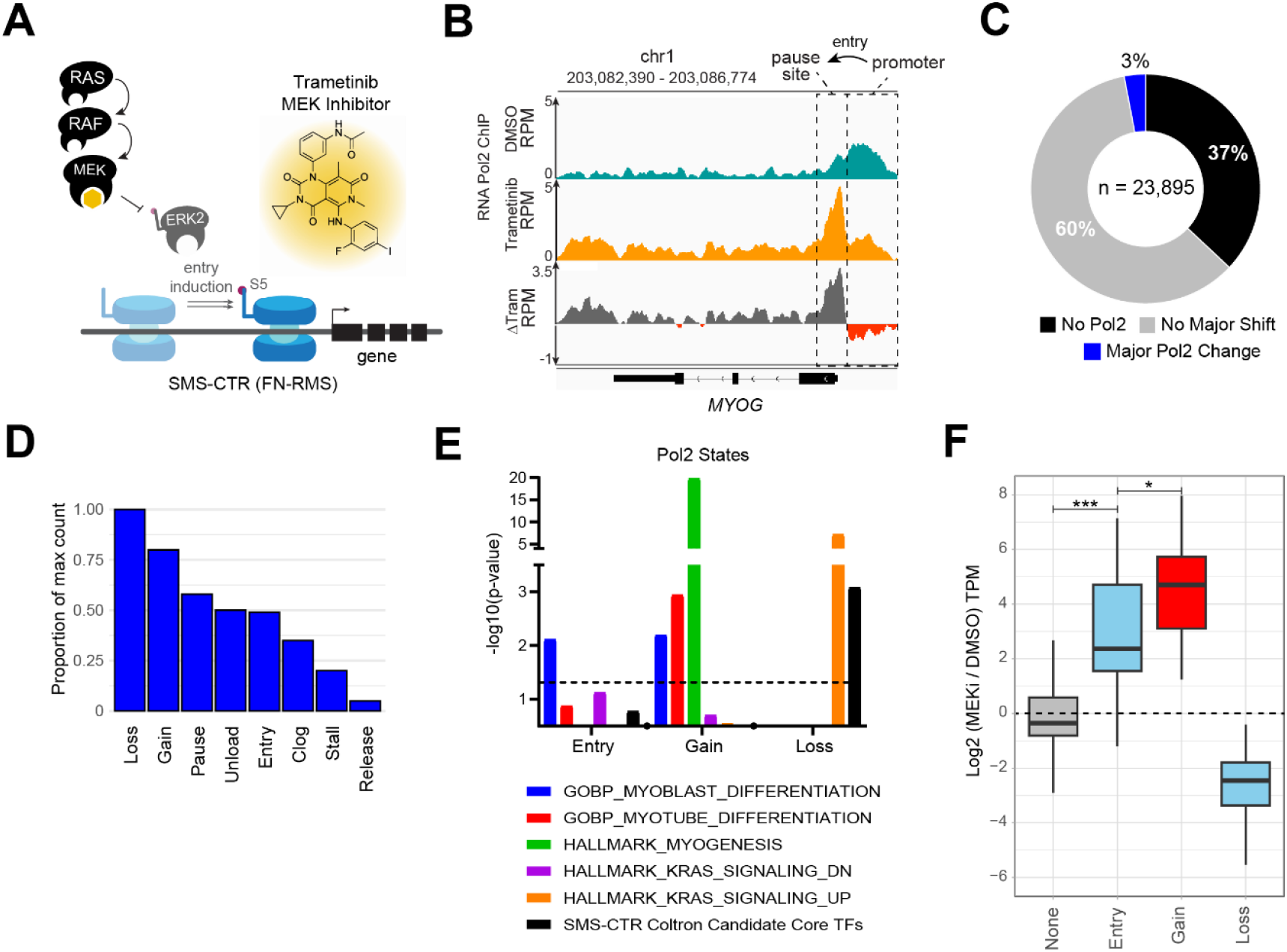
MEK pathway inhibition improves Pol2 recruitment and processivity in FN-RMS. **A**. Model of the described impact of trametinib in SMS-CTR fusion-negative rhabdomyosarcoma cells. **B**. Genome browser tracks of Pol2 ChIP-seq data showing Pol2 *Entry*, a gain in Pol2 at the TSS region and accompanying loss in the promoter region, at the *MYOG* locus. **C**. Donut plot showing percentage of genes showing major Pol2 perturbation, no major change and not considered due to low Pol2. **D**. Bar plots showing the proportion of perturbed genes in each Pol2 state relative to the most frequent state. **E**. Bar plot showing the enrichment of genes in Pol2 final states for known functional sets of genes. Dashed line shows the corresponding cutoff for p values < 0.05. Dots along the x-axis separate tranches of bars for each Pol2 state. **F**. Boxplots showing the distribution of L2FC gene expression values for genes in unperturbed, Pol2 entry, gain and loss states where the difference between basal and trametinib treated state was >= 10 TPM. Unperturbed genes are in grey, Pol2 states which reflect a gain in a preceding region are in red, and other Pol2 states are in blue. The differences are statistically significant as determined from a Mann-Whitney U-test.

Applying compPASS to the data from this study, genome-wide evaluation revealed that major Pol2 changes are rare, occurring in just 3% of genes (Fig. 4C). Unlike in the previous, canonical *Pausing* case, no single mode dominated (Fig. 4D, Supplementary Fig. 4A). *Pol2 Loss* is the most abundant mode (Supplementary Fig. 4B), with genes in this mode showing a strong drop along the entire gene region (Supplementary Fig. 4C). Gene set enrichment analysis reveals that, while genes labeled *Entry* are enriched for myoblast differentiation (including *MYOG*), the heretofore unmeasured *Pol2 Gain* category is heavily enriched for myoblast differentiation, myotube differentiation, and hallmark myogenesis (Fig. 4E, Supplementary Fig. 4D).Meanwhile, *Pol2 Loss* is enriched for genes in hallmark KRAS signaling up. Genes showing *Pol2 Gain* show increased gene expression after trametinib treatment, even more so than Pol2 entry, while genes showing *Pol2 Loss* show the opposite (Fig. 4F). This aligns with their finding that MYOG *Entry* and transcriptional upregulation leads to activation of its downstream targets and thereby induces differentiation. The compPASS analysis approach thus highlights that a rare mode of Pol2 regulation (a switch from Stalling to Entry) is a unique epigenetic lynchpin in the cancer-causing differentiation block that defines RAS-driven RMS.

### compPASS shows non-pausing Pol2 aggregation gene classes in FP-RMS

In the fusion-positive rhabdomyosarcoma cell-line RH4, a core cancer driving protein is the PAX3::FOXO1 fusion transcription factor (TF) which binds chromatin and maintains the expression of other core regulatory TFs and downstream target genes. Dual degradation of p300/CBP with dCBP1 induces a specific and substantial loss of P3F-regulated genes (Fig. 5A). While Pol2 binding sites showed increases and decreases in Pol2 abundance in near equal number following d-CBP1 treatment in our prior work^9^, when we stratified Pol2 status at downregulated genes, we saw that Pol2 was gained at genes which overlapped a CpG island (Fig. 5B). We compare here the difference in Pol2 activity state change at *PIPOX* and *MYOD1*, two highly expressed genes and core biomarkers for FP-RMS growth (Fig. 5C). Previously, we defined this Pol2 accumulation at CpG-overlapping genes and noted how it defies the typical *Pausing* paradigm.

**Figure 5:**
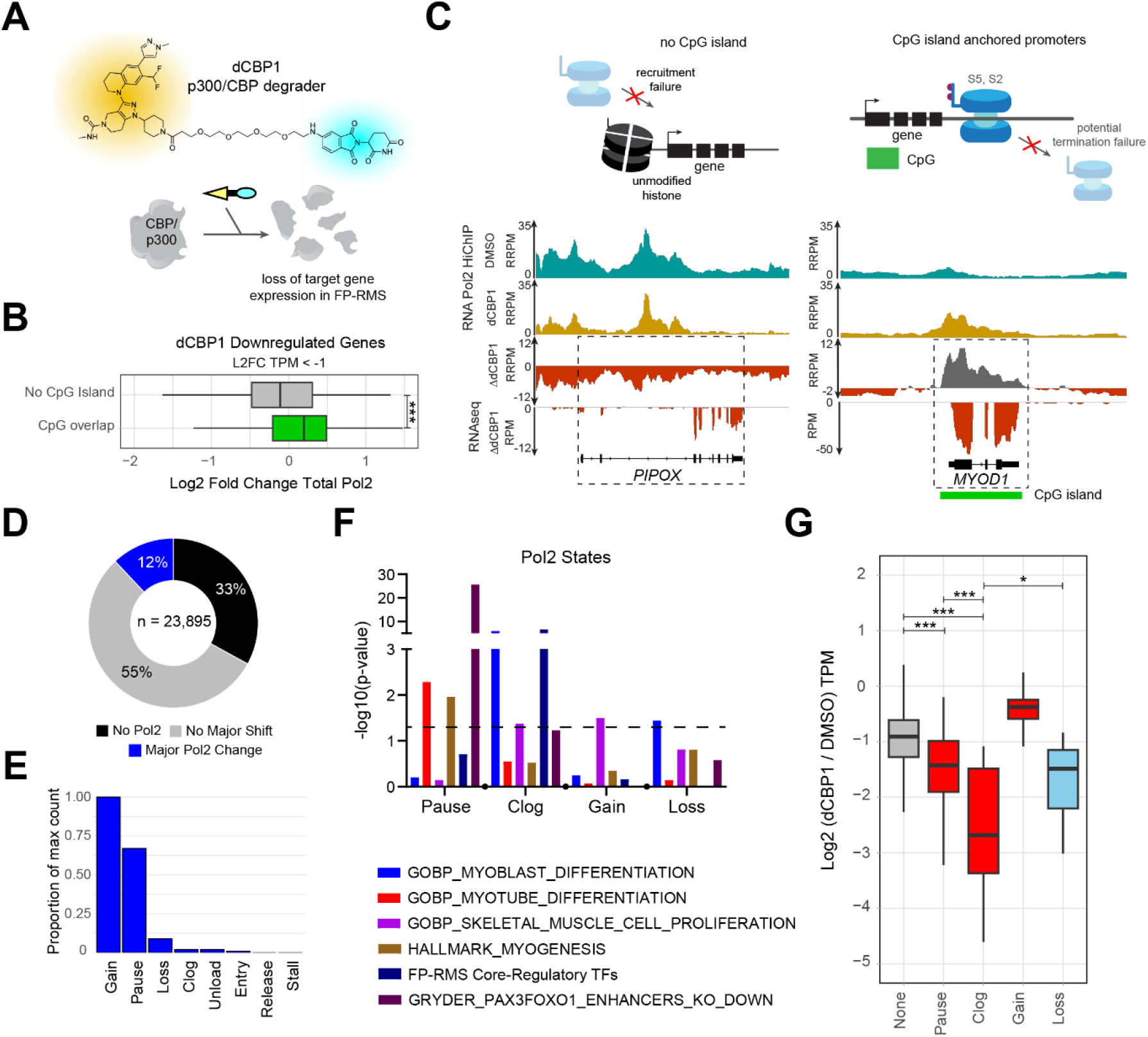
p300/CBP degradation causes rare clogging events that impede cancer growth in FP-RMS. **A**. Model depicting structure of d-CBP1 and degradation of CREBBP and EP300. **B**. Box plot showing the distribution of log2 fold change total Pol2 between downregulated genes do and do not overlap a CpG island. The difference is statistically significant as determined from a Mann-Whitney U-test. **C**. Genome browser view of Pol2 HiChIP across DMSO and d-CBP1 treatment at the PIPOX and MYOD1 loci (bottom) and accompanying proposed models of drug impact at gene-proximal regions (top). **D**. Donut plot of the percentage of genes highly perturbed, unshifted, and with trivial Pol2. **E**. Bar plots showing the proportion of perturbed genes in each Pol2 state relative to the most frequent state. **F**. Bar plot showing the enrichment of genes in Pol2 final states for known functional sets of genes. Dashed line shows the corresponding cutoff for p values < 0.05. Dots along the x-axis separate tranches of bars for each Pol2 state. **G**. Boxplots showing the distribution of L2FC gene expression values for genes in unperturbed, Pol2 entry, gain and loss states where the difference between basal and d-CBP1 treated state was >= 10 TPM. Unperturbed genes are in grey, Pol2 states which reflect a gain in a preceding region are in red, and other Pol2 states are in blue. The differences are statistically significant as determined from a Mann-Whitney U-test.

Genome-wide, only about 12% of genes showed a major Pol2 change after dCBP1 (Fig. 5D). Most genes are classified as experiencing *Pol2 Gain, Pol2 Loss* and *Pausing* (Fig. 5E, Supplementary Fig. 5A, Supplementary Fig. 5B). *Pol2 Gain* is far more common than *Clogging* and shows greater maximum gain at the TSS (Supplementary Fig. 5C). However, *Clogging* and *Pausing* genes are far more enriched for biologically relevant genes, like those regulated by P3F and FP-RMS core-regulatory transcription factors (Fig. 5F, Supplementary Fig. 5D). Looking at the transcription, genes which experience Pol2 clogging show greater downregulation than the other major categories for this perturbation (Fig. 5G). Despite containing almost 50x as many genes as the *Clogging* category, the *Pol2 Gain* category does not capture the most impacted genes or the most cell-state relevant genes in this context.

## DISCUSSION

compPASS is an open access pipeline for quantifying genes undergoing Pol2 state shifts and identifying which steps in the transcriptional life cycle are being influenced by any given chemical or genetic perturbation. To our knowledge, this tool and accompanying work are the first to try to capture Pol2 activity shifts genome-wide and provide definitions of gene-sets from the changes across four major regions relative to the promoter, TSS, gene body and TES of gene loci. We also provide distinct definitions of the stalling, pausing and termination ratios which contextualize Pol2 disruption at steps of initiation, pause release and termination.

Previous efforts to assess genes defined de novo as paused, like in Day et al^5^, used a loose cutoff of pause ratio > 2 and, in human cells, identified more than 30% of genes as paused. In L3.6pl cells, the same approach would label 87% (13,771) of the considered genes paused before CDK9 inhibition. Additionally, most (4,519 of 4,620) of the genes classified as *Pausing* here would have been called paused in the basal state by this metric (see Supplementary Table 2). While the regions used differ (their gene body site extends 3 kb past the TES), a much larger gene set weakens the resulting paused gene set’s enrichment for known ontologies.

We saw chemical perturbations with broader epigenetic impacts show a lower percentage of strongly perturbed genes, between 2% and 12% of our overall set of 23,895 genes as compared to 30% after CDK9 inhibition. The percentage of depleted genes across cell lines remained between 30% and 40%, suggesting that the driver of this difference was the kinds of perturbations taking place.

The most striking finding from compPASS is that biology hides in the rare modes.After p300/CBP degradation in fusion-positive rhabdomyosarcoma, only ∼2% of Pol2 perturbed genes are classified as clogging - yet that 2% is enriched for PAX3-FOXO1 targets and core-regulatory transcription factors and shows greater downregulation than any other class. The fifty-fold larger *Pol2 Gain* mode does not. The number of core regulatory genes classified as *Clogging* may reflect a structural vulnerability. Without essential co-activators like p300 to maintain the condensate, Pol2 may fail to biochemically switch into the productive elongation and splicing compartments, leading to terminal arrest within the gene body^11,12^. This also aligns with the loss of interactions with spliceosome factors seen when PAX3::FOXO1 is mutated and no longer able to colocalize with p300 (ref ^9^); Pol2 cluster collapse may also be accompanied by PAX3::FOXO1’s exclusion from transcriptionally active condensates.

The compPASS tool’s findings are partially dependent on the gene reference provided. This is user-modifiable within the pipeline but, even with isoform selection, the human genome contains varied gene contexts that could impact Pol2 positioning. This study does not exclude genes with diverging TSS, converging TES, or overlapping genes from the results, as prior testing showed that most gene sets showed minimal bias for specific modes. It is unclear to what extent gene contexts align with known classes of genes, i.e. many housekeeping genes are in dense gene neighborhoods^13^. In lieu of more complex signal deconvolution techniques to parse contributing ChIP-style signals across hundreds of overlapping gene loci, user-directed gene reference choice offers a practical way to mitigate the impact of ambiguous Pol2 signal.

We also sought to simplify the classification of Pol2 modes by excluding mixed modes, i.e. genes experiencing simultaneous Pol2 entry and pausing. Manual curation of sets of mixed-mode cases identified genes with low read coverage such that fold-change metrics overestimated the importance of the perturbation. Therefore, genes are preferentially assigned to discrete classes in this version of compPASS. Prior studies clarify the difference in Pol2 positioning along the gene body by targeting Serine 5 and Serine 2 phosphorylated Pol2 with ChIP-seq^14,15^. However, adopting this strategy here would limit the tool’s ability to assess Pol2 changes within the gene body, given the promoter specificity of Ser5P and the restricted coverage and antibody-specificity issues associated with Ser2P (ref ^16^). Further study, likely targeting multiple Pol2 modifications in parallel, would be required to clarify if mixed Pol2 shift modes represent genuine biology or rare artifacts found at dense gene neighborhoods. Even so, restricting genes to singular Pol2 state shifts still highlights biologically critical genes through their distinct response signature.

### Limitations and Future Directions

RNA Pol2 driven transcription has many biochemically diverse states, diverse 3D organizational modes, and diverse bottlenecks of regulation. While powerful, the current view of Pol2 transitions is restricted to considering (1) Pol2 movement in 1D, ignoring 3D chromatin modes, (2) makes assumptions based on bulk data, averaging millions of cells in a variety of epigenetic states, and (3) focuses on total Pol2 signal, with no consideration for various Pol2 phosphorylation states. In real biology, 1D transcription and 3D architectural processes are deeply intertwined and often antagonistic. For example, high Pol2 engagement can coordinate multiple enhancer-promoter interactions within strong condensates, but Pol2 placement can restrict the placement of insulating boundaries by cohesin and CTCF^17^. Integrating our 1D Pol2 metrics with 3D models will be critical for distinguishing different types of clusters and corresponding 3D context specific Pol2 activity shifts. Current models suggest that transcription activation relies on the interaction radius of phase-separated enhancer-promoter condensates. Enhancer-driven clusters require large interaction radii and dynamic co-activator recruitment, making them highly vulnerable to collapse upon targeted degradation^18^. Stratifying Pol2 activity state shifts by cluster size and differential occupancy of cofactors may be key to extending sequencing-based analyses of Pol2 to questions of condensate occupancy.

Future single-molecule imaging and multi-omic approaches will be required to define how these 1D state shifts correspond to the assembly, dynamic exchange, and collapse of 3D biomolecular condensates. Live cell imaging approaches of transcription have the benefit of highly resolved temporal dynamics but are generally restricted to single genes and lack genomic positional information. To bridge the gap between throughput and resolution, we envision that, with improvements in single-cell ChIP-seq for Pol2 (or possibly single-cell nascent RNA measurements such as NET-seq^7^/ChRO-seq^19^), future work may overcome some of these limitations. In this current work, we did not attempt to benchmark compPASS compatibility with nascent RNA sequencing data, but the software is entirely compatible with these datatypes and may only require a close evaluation of changes to the thresholds that define categories of Pol2 state shifts. We also envision that a more complete and mechanistically precise assay for capturing transcriptional regulation will allow the field to venture past these current conceptual and technological barriers.

Broader use of this tool will enable deeper understanding of this understudied element of epigenetic regulation and help move the field beyond a myopic focus on Pol2 pausing alone.

## STAR METHODS

### Key resources table

| REAGENT or RESOURCE | SOURCE | IDENTIFIER |
| --- | --- | --- |
| **Deposited data** |  |  |
| GSE208146 | Asante et al,<sup>9</sup> | N/A |
| GSE319181 | Khan et al,<sup>20</sup> | N/A |
| GSE116344 | Gryder et al,<sup>21</sup> | N/A |
| GSE260915 | Chin et al,<sup>10</sup> | N/A |
| GSE50622 | Cayrol et al,<sup>22</sup> | N/A |
| GSE271732 | Ekstrom et al,<sup>23</sup> | N/A |
| GSE85169 | Yohe et al,<sup>8</sup> | N/A |
| **Software and algorithms** |  |  |
| R, v4.4.2 | R Core Team | https://www.r-project.org/ |
| Bedtools, v2.30.0 | Quinlan et al,<sup>24</sup> | https://doi.org/10.1093/bioinformatics/btq033 |
| Samtools, v1.19.2 | Li et al,<sup>25</sup> | https://doi.org/10.1093/bioinformatics/btp352 |
| Deeptools, v3.5.5 | Ramirez et al,<sup>26</sup> | http://doi.org/10.1093/nar/gkw257 |
| Python version 3.8 | Python Software Foundation | https://www.python.org/ |
| compPASS pipeline | This paper | https://github.com/GryderLab/compPASS |

## METHOD DETAILS

### Inputs for Pipeline

The required inputs for compPASS are a control and case/perturbed BAM files of RNA Pol2 ChIP-seq or HiChIP and their accompanying index files (.bam.bai). For proper scaling, it is also necessary to know the total number of aligned reads or, if employing spike-in, the total number of aligned orthogonal species reads which are used in the command-line of the script. The pipeline allows the user to provide their own gene reference (in BED format) but one is provided, derived from the RefSeq hg38 set of genes.

### Regions of Interest

We define four regions of interest at gene loci, relative to the TSS and TES: promoter region (-800 to -30 from the TSS), TSS region (-30 to +300 from the TSS), gene body region (+300 to TES) and TES region (TES to +4000 from TES). These align broadly with regions denoted in prior studies.

### Selection of Genes for Consideration

To avoid overlapping regions of interest, we remove genes from consideration that are less than 2,000 bp from TSS to TES (removes 5,000 genes, n = 23,895 genes remaining). Additionally, to simplify the application of the tool across cells derived from tissues of either sex, we exclude genes on chromosome Y. Downstream filtering occurs at runtime, in which genes with promoter-proximal (TSS region) Pol2-associated reads in the bottom 30th percentile (virtually no Pol2 bound at promoter region or TSS region) are excluded from the analysis set.

### Gene Categorization by Intrinsic Properties

Prior to ratio-based classification, genes are categorized as coding, housekeeping, or transcription factors according to prior annotation from ENCODE, with the full list used here provided in Supplementary Table 1. Additionally, genes are categorized according to length (3 bins, < 10Kb, 10 – 100Kb and > 100Kb where kb means kilobase-pairs), whether they overlap a CpG island, and whether they experience a gene overlapping event (i.e. diverging promoter regions, colliding TES regions, sense/anti-sense promoter region overlap, sense/antisense TES region overlap and/or direct gene body overlap). While enrichments provided by the pipeline use this provided gene table, users may provide their own comparable table in tab-separated value format, provided that the categorizations are binary (TRUE or FALSE) for each column.

### Calculation of Gene Pausing, Stalling and Termination Ratios

We define each ratio by taking the normalized read density of the preceding region and dividing it by the normalized read density of the subsequent region. Read counts are captured via bedtools (version 2.30.0, ref ^24^). Normalization is in reads per million, RPM = read_count * (1,000,000 / total_aligned_reads) and uses samtools (version 1.19.2, ref ^25^). Therefore, the Stalling Ratio is defined using the promoter and TSS regions as (Normalized Promoter Region RPM / Length of Promoter Region) / (Normalized TSS Region RPM / Length of TSS Region). If you replace the former and latter values, the equation is the same for the Pausing Ratio (TSS region and gene body region) and Termination Ratio (gene body region and TES region).

### Gene Classification by Ratio

Initial gene classification is done as follows: if the log2 fold change (L2FC) between case and control of the first region is greater than 0.799 and the L2FC between case and control at the second region is less than 0, while the ratio L2FC is greater than 0.999 that region and the final ratio is greater than or equal to 2, the gene is considered to be experiencing that class of activity state shift. For example, in the case of pausing, if the L2FC of read density in the TSSR is greater than 0.799, the L2FC of read density in the gene body is less than 0, the L2FC pausing ratio is greater than 0.999 and the resultant ratio is greater than or equal to 2, the gene is *Pausing*. The inverse is true for releasing, such that no *Pausing* gene can be considered *Releasing*. However, genes can experience multiple modes according to the initial metric. Genes are classified based on ratio with the greatest magnitude, with the class dependent on the sign. For example, a gene which meets the criteria for *Stalling* and *Pausing*, with a L2FC of the Stalling Ratio of 1.23 and a Pausing Ratio of 1.62 would be classified as *Pausing*.

### Profile Plots by Gene Class

We use deeptools (version 3.5.5, ref ^26^) to plot the average Pol2 ChIP-seq abundance for each set of classified genes across a window of -3,000 bp from the TSS to +4,000 bp from the TES. The inputs for computeMatrix are generated with bamCoverage at a bin size of 10 bp and are normalized using the provided scale factors.

### Outputs of compPASS

For each successful run of the compPASS pipeline, the user should receive:

- A tab-separated values table which captures the RPM values for sample A’s regions of interest and calculated ratios, RPM values for sample B and calculated ratios and the log2 of sample B’s RPM / sample A’s values (representing the L2FC)
- Scatterplots showing the L2FC of the following paired regions – promoter region and TSS region, TSS region and gene body region, and gene body region and TES region. These plots also highlight in red or blue respectively, the genes classified as stalling or entering, pausing or releasing and clogging or unloading.
- Radar plots showing the number of genes assigned to each of the eight classifications
- Text files listing genes in each class
- Average plots of Pol2-associated RPM signal across the loci of genes in each class
- Enrichment dot plots for the provided sets in the gene categories tables

### Statistical analysis

Statistical tests were performed using the two-sample wilcox function (Mann-Whitney U-test) in R (version 4.4.2).

## Data availability

The compPASS pipeline was built with R and Python; the tool and dependencies are available at https://github.com/GryderLab/compPASS.

## ACKNOWLEDGEMENTS

We would like to thank Issra Osman and Diana Chin for their prior experimental work on RH4 RNA Pol2 HiChIP which helped demonstrate the landscape of Pol2 failure modes in subsequent analysis. This work made use of the High-Performance Computing Resource in the Core Facility for Advanced Research Computing at Case Western Reserve University.

## AUTHOR CONTRIBUTIONS

Yaw Asante [Methodology, Investigation, Data curation, Formal Analysis, Software, Visualization, Writing-original draft]; Berkley Gryder [Conceptualization, Supervision, Visualization, Writing-revisions, Final approval]

