## Supplementary Figures S1-5 for "Rare RNA Polymerase II failure modes mark the cancer-driving genes most affected by epigenetic perturbation"

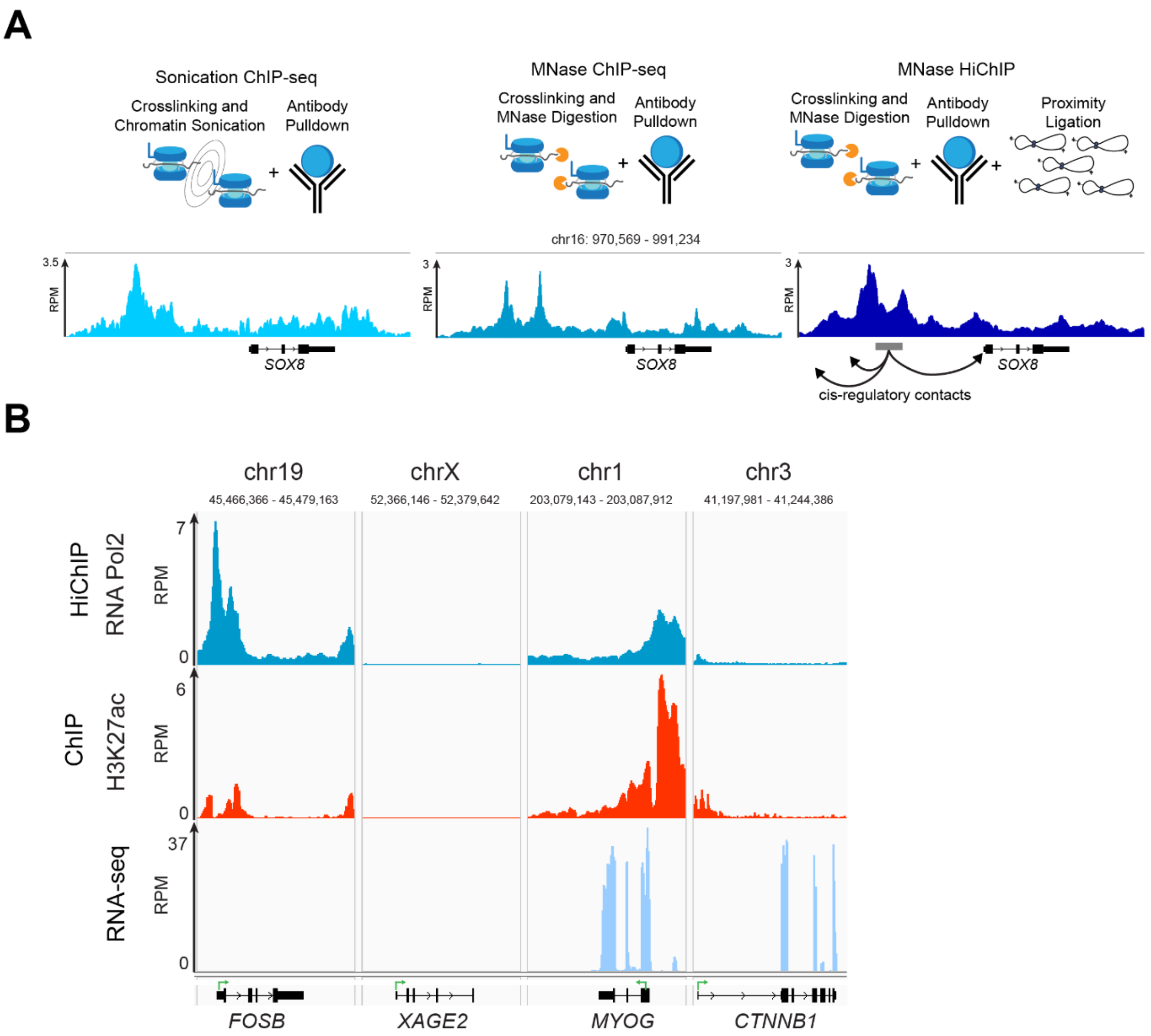


**Figure S1: RNA Polymerase 2 profile capture can reflect gene activity but accumulation alters expectations.**

**A.** Different types of ChIP techniques used to capture RNA Pol2 positioning (top) and corresponding genome browser track views of the Pol2 reads at the *SOX8* locus (bottom). HiChIP captures long-range contacts as well, as noted in the MNase HiChIP example.

**B.** Genome browser track views of RNA Pol2 HiChIP, histone acetylation (H3K27ac) and gene expression (RNA-seq) at four loci which show conditions of high Pol2-low expression, no Pol2-no expression, high Pol2-high expression, and low Pol2-high expression. Sense directions of each gene are noted with the green arrows.

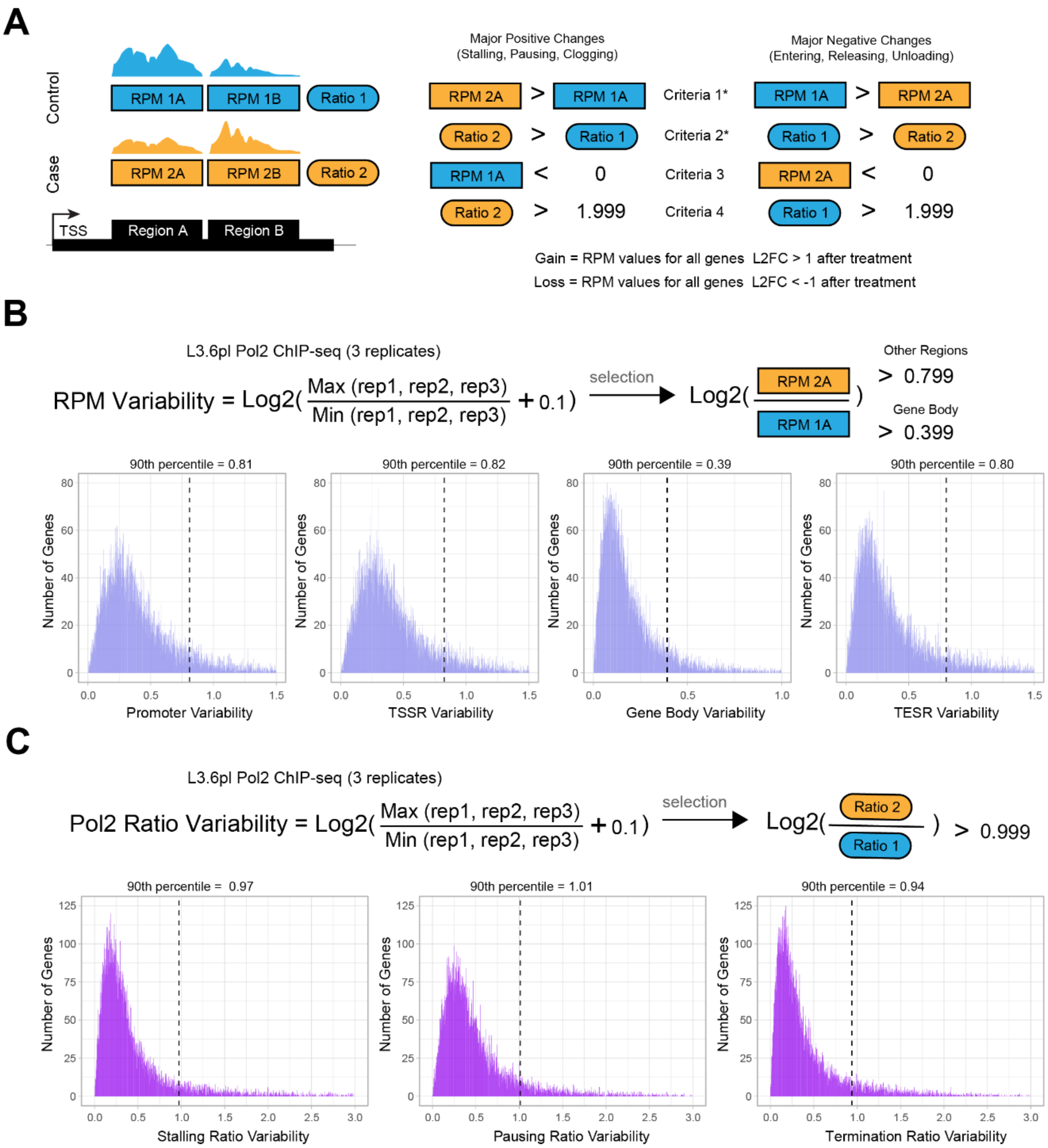


**Figure S2: Approach to capturing substantial changes in RNA Pol2 ratio and abundances at gene-proximal regions**

**A.** General framework and four criteria for assigning the eight perturbed Pol2 states

**B.** Formula for Pol2 variability for Pol2 abundance across three replicates of L3.6pl Pol2 ChIP-seq (top) and histograms of Pol2 variability across the four gene-proximal regions (bottom). The assigned cutoffs for each region are selected from the approximates of the histogram 90$^{th}$ percentile cutoff.

**C.** Formula for Pol2 variability from the three Pol2 ratios across three replicates of L3.6pL Pol2 ChIP-seq (top) and histograms of Pol2 variability across stalling, pausing and clogging ratios (bottom). The assigned ratio cutoffs for each region are selected from the approximates of the histogram 90$^{th}$ percentile cutoff.

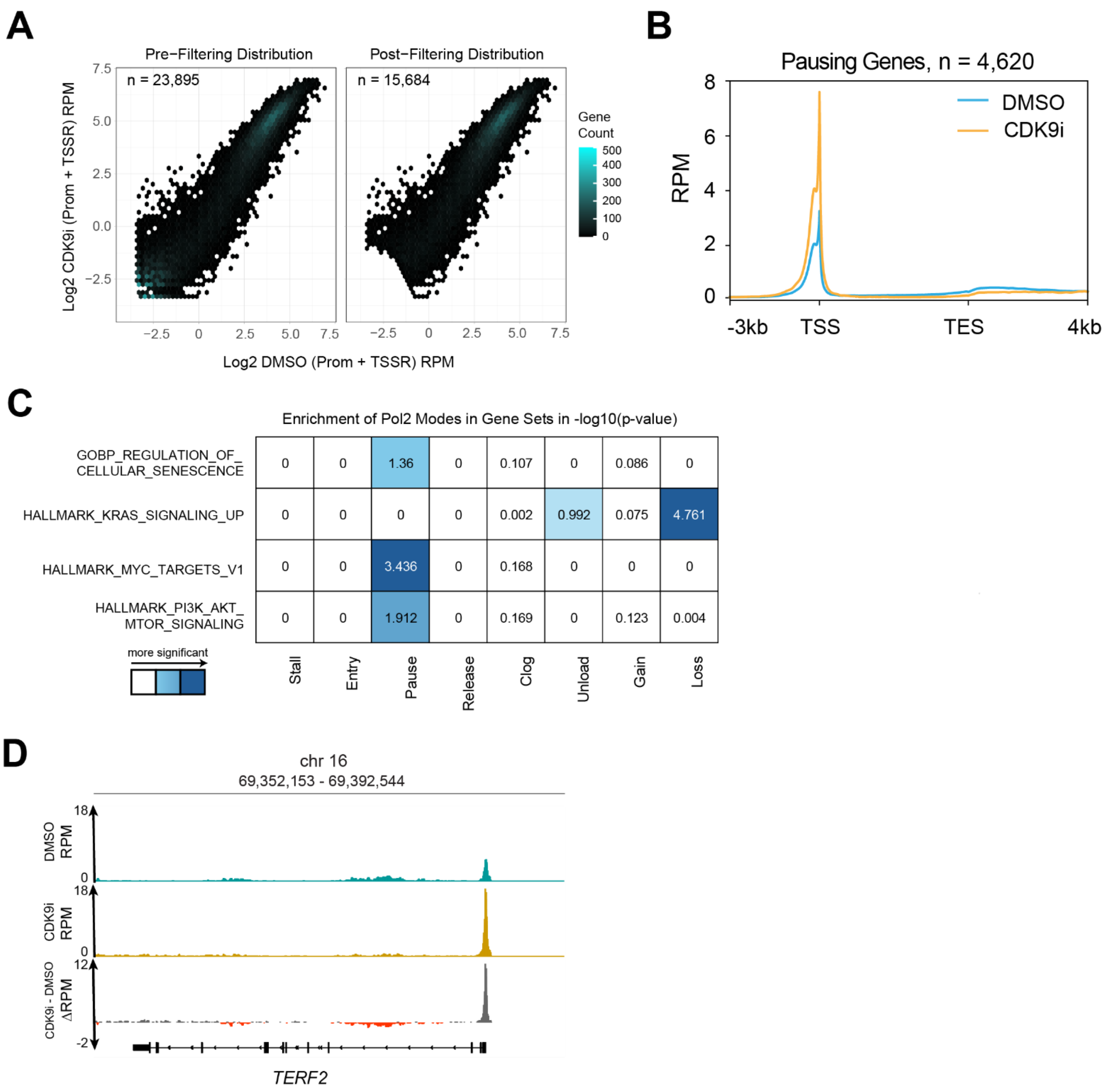


**Figure S3: CDK9 inhibition causes pausing and loss at oncogenic genes in PDAC.**

**A.** Hex plots showing the distribution of genes by combined promoter and TSS across control (DMSO-treated, x-axis) and perturbed (enitociclib-treated, y-axis). Gene counts noted before and after filtering.

**B.** Metagene plot of Pol2 ChIP-seq from all genes categorized as pausing between DMSO and enitociclib treatment.

**C.** Matrix of enrichment results from all categorized genes and select relevant sets which show significant enrichment in at least one category. White text indicates highly significant enrichment values (>= 3.3) from a hyper-geometric test.

**D.** Genome browser view of an TERF2, an example negative cell senescence regulator, showing pausing after CDK9 inhibition.

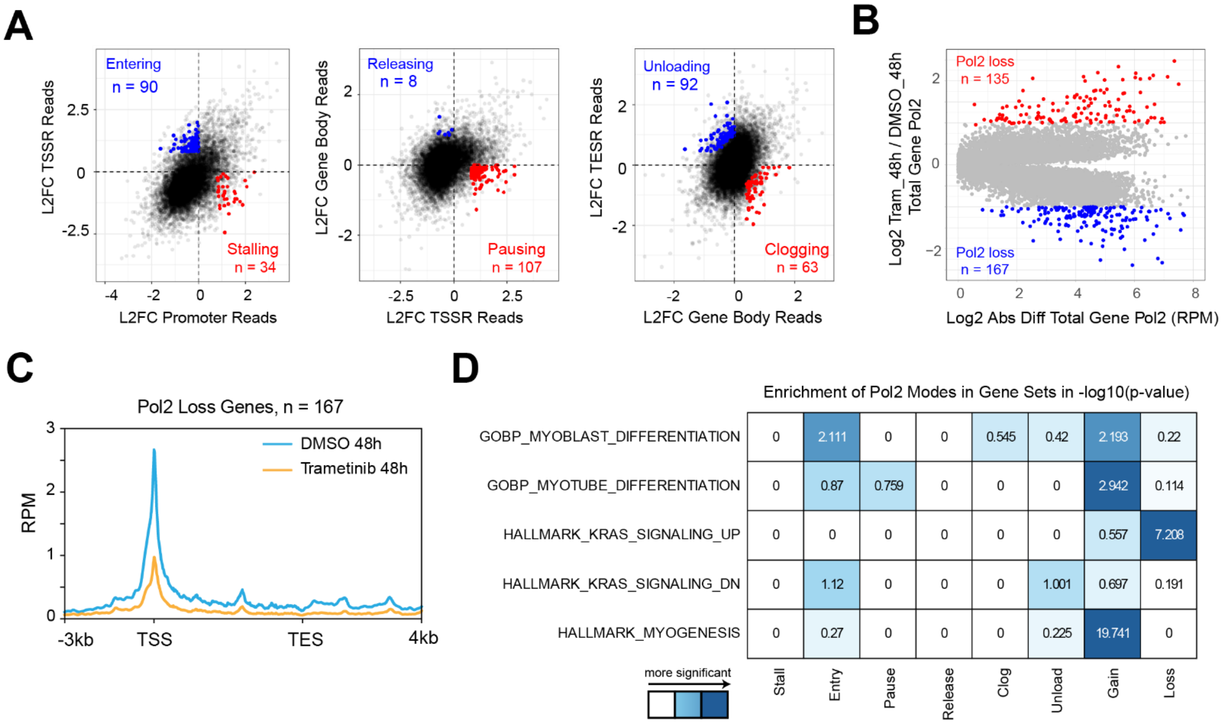


| | Stall | Entry | Pause | Release | Clog | Unload | Gain | Loss |
|---|---|---|---|---|---|---|---|---|
| GOBP_MYOBLAST_DIFFERENTIATION | 0 | 2.111 | 0 | 0 | 0.545 | 0.42 | 2.193 | 0.22 |
| GOBP_MYOTUBE_DIFFERENTIATION | 0 | 0.87 | 0.759 | 0 | 0 | 0 | 2.942 | 0.114 |
| HALLMARK_KRAS_SIGNALING_UP | 0 | 0 | 0 | 0 | 0 | 0 | 0.557 | 7.208 |
| HALLMARK_KRAS_SIGNALING_DN | 0 | 1.12 | 0 | 0 | 0 | 1.001 | 0.697 | 0.191 |
| HALLMARK_MYOGENESIS | 0 | 0.27 | 0 | 0 | 0 | 0.225 | 19.741 | 0 |

**Figure S4: MEK inhibition leads to broad impacts in Pol2 positioning including acute gain at muscle lineage genes.**

**A.** Scatterplots showing genes in each state based on the distribution of fold change values from their respective gene-proximal regions. Pol2 states reflecting a gain in the preceding region (Stalling, Pausing, Clogging) in red, Pol2 states reflecting a loss in the preceding region in blue.

**B.** Scatterplot showing the log2 absolute difference of total Pol2 against the fold change of total Pol2. Total Pol2 is the sum of Pol2 at all gene-proximal regions. Pol2 gain genes in red, Pol2 loss genes in blue.

**C.** Metagene plot of Pol2 ChIP-seq from all genes categorized as Pol2 loss between DMSO and trametinib treatment.

**D.** Matrix of enrichment results from all categorized genes and select relevant sets which show significant enrichment in at least one category. White text indicates highly significant enrichment values (>= 3.3) from a hyper-geometric test.

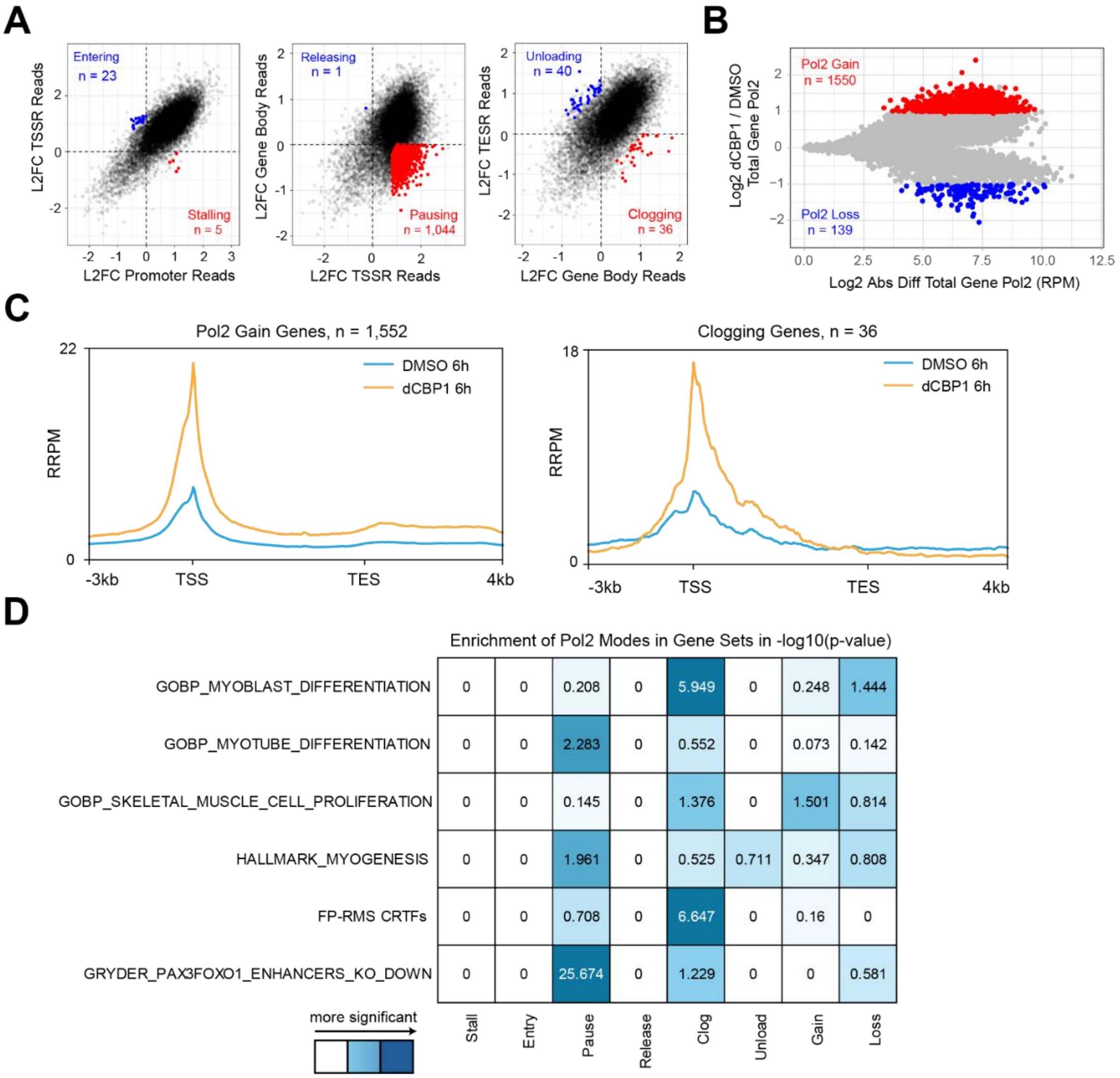


**Figure S5: d-CBP1 induces distinct Pol2 accumulation pattern that is enriched for core-regulatory transcription factors.**

**A**. Scatterplots showing genes in each state based on the distribution of fold change values from their respective gene-proximal regions. Pol2 states reflecting a gain in the preceding region (Stalling, Pausing, Clogging) in red, Pol2 states reflecting a loss in the preceding region in blue.

**B**. Scatterplot showing the log2 absolute difference of total Pol2 against the fold change of total Pol2. Total Pol2 is the sum of Pol2 at all gene-proximal regions. Pol2 gain genes in red, Pol2 loss genes in blue.

**C**. Metagene plot of Pol2 ChIP-seq from all genes categorized as Pol2 gain and Pol2 clog between DMSO and d-CBP1 treatment.

**D**. Matrix of enrichment results from all categorized genes and select relevant sets which show significant enrichment in at least one category. White text indicates highly significant enrichment values (>= 3.3) from a hyper-geometric test.
